# Comparing neural models using their perceptual discriminability predictions

**DOI:** 10.1101/2023.11.17.567604

**Authors:** Jing Yang Zhou, Chanwoo Chun, Ajay Subramanian, Eero P. Simoncelli

**Affiliations:** Flatiron Institute, New York, NY 10010; Weill Cornell Medical College, New York, NY 10065; New York University, New York, NY 10003

## Abstract

Internal representations are not uniquely identifiable from perceptual measurements: different representations can generate identical perceptual predictions, and similar representations may predict dissimilar percepts. Here, we generalize a previous method (“Eigendistortions” – Berardino et al., 2017) to enable comparison of models based on their metric tensors, which can be verified perceptually. Metric tensors characterize sensitivity to stimulus perturbations, reflecting both the geometric and stochastic properties of the representation, and providing an explicit prediction of perceptual discriminability. Brute force comparison of model-predicted metric tensors would require estimation of human perceptual thresholds along an infeasibly large set of stimulus directions. To circumvent this “perceptual curse of dimensionality”, we compute and measure discrimination capabilities for a small set of most-informative perturbations, reducing the measurement cost from thousands of hours (a conservative estimate) to a single trial. We show that this single measurement, made for a variety of different test stimuli, is sufficient to differentiate models, select models that better match human perception, or generate new models that combine the advantages of existing models. We demonstrate the power of this method in comparison of (1) two models for trichromatic color representation, with differing internal noise; and (2) two autoencoders trained with different regularizers.

## 1 Introduction

Stimulus discriminability is the most reliable, widely-used, and well-understood measurement of perception. Discriminability is believed to reflect the degree of change in neural responses induced by small stimulus perturbations [1, 2, 3], and methods for estimating perceptual discrimination are highly refined and efficient. In this paper, we propose an efficient method to compare neural models based on their ability to account for human perceptual discriminability.

Many existing methods for model comparison assess the similarity between neural representations of stimuli [5, 6, 7, 8, 9, 10]. This approach is grounded in the belief that models that have similar stimulus representations are similar. However, similarity identified by such methods may not translate into similarity between the models’ perceptual predictions. In Figure 1, we show representations of 3 models, each describing stochastic responses of two neurons to a one-dimensional stimulus (Fig. 1A). The first two representations are very different both visually and according to several quantitative metrics (Fig. 1A), but they share the same perceptual discriminability predictions. Conversely, the first and third representations are visually similar and are judged as highly similar by current metrics (Fig. 1C), but their predicted perceptual discriminabilities significantly differ. To resolve this issue, we summarize a model’s predicted discriminability using a metric tensor, a positive semi-definite matrix that characterizes the model’s sensitivity to stimulus perturbations, and propose to compare models by their ability to predict human perceptual discriminability. Characterizing human discriminability for a high-dimensional stimulus (e.g., an image) requires measuring thresholds over a huge number of dimensions, and would need thousands of hours of human subject time. Our method circumvents this “perceptual curse of dimensionality” by requiring human judgements only along the stimulus perturbation dimensions that best differentiate the two models being compared.

**Figure 1:**
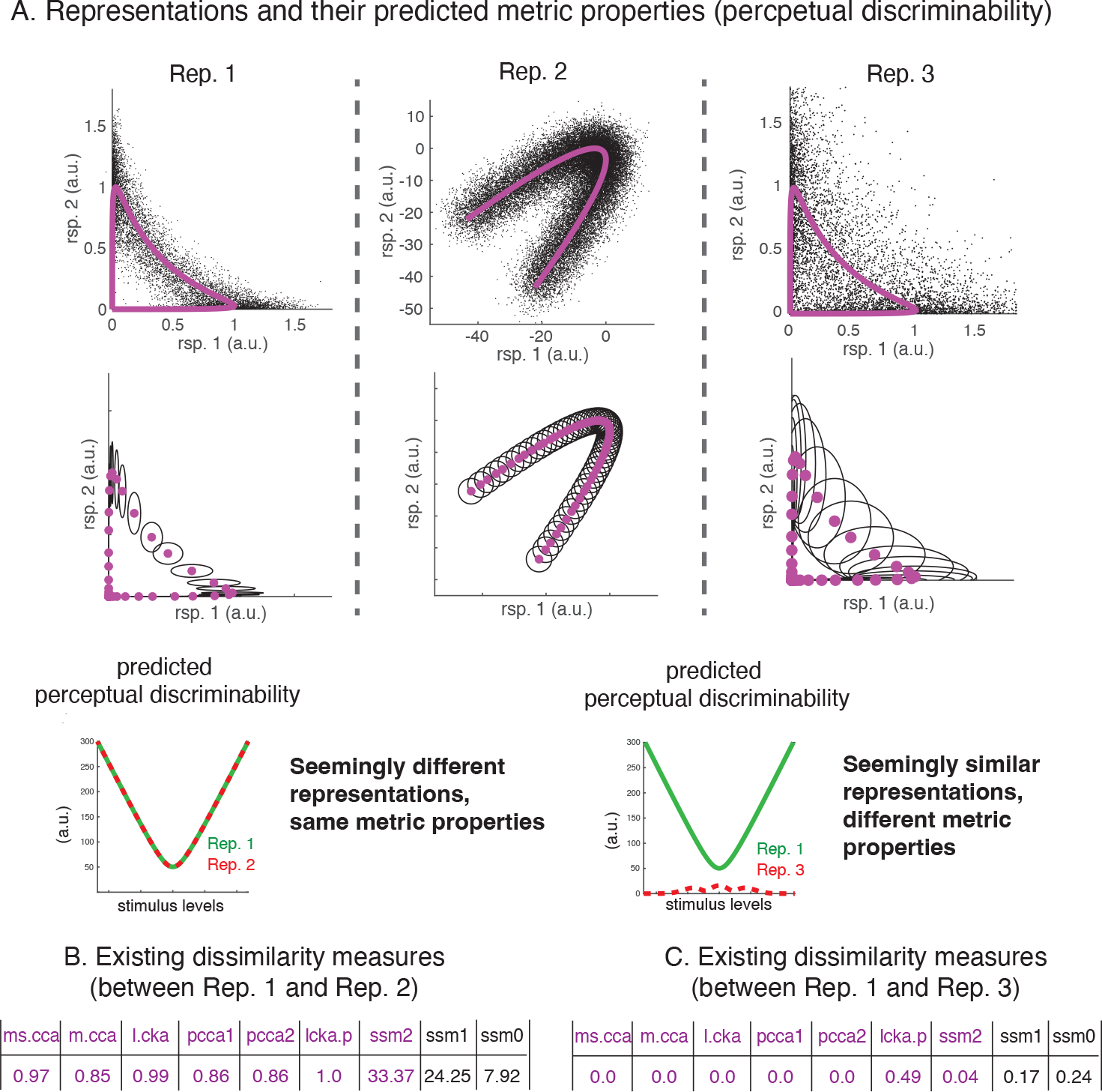
Existing model comparison methods do not reflect similarity of perceptual discriminability predictions. **A**. Responses of three stochastic neural representations (Rep. 1, Rep. 2, Rep. 3). Each plot depicts two neurons’ stochastic responses (black points), and mean responses (magenta) for a family of stimuli parameterized by a one-dimensional parameter. Rep. 1 and Rep. 2 produce identical perceptual discriminability along the stimulus axis, even though their mean responses are quite different. Rep. 1 and Rep. 3 have the same mean responses, but their predicted perceptual discriminabilities along the stimulus axis are very different. Second row: discretely sampled neural representations, illustrating mean and covariance of responses to selected stimuli. Third row: predicted perceptual discriminability for each representation, along the stimulus axis. **B**. Comparisons of Rep.1 and Rep.2 yield large dissimilarity scores for a set of comparison models. **C**. Comparisons of Rep. 1 and Rep. 3 yield small dissimilarity scores for all model. Included models are: (1) CCA with mean squared correlation coefficient; (2) CCA with mean correlation coefficient; (3) linear CKA; (4) projection-weighted CCA projected using the first representation’s weights; (5) projection-weighted CCA using the second representation’s weights; (6)linear CKA prime distance; (7) stochastic shape metric using only mean responses (equivalently, Procrustes shape metric on mean responses); (8) stochastic shape metric using both mean and covariances; and (9) stochastic shape metric using only covariances. For all CCA-related metrics, we subtract the standard metric from one to estimate dissimilarity. The first 7 methods were summarized in Ding et al. 2021, and the last 3 metrics were summarized in [4].

In summary, this article makes the following contributions:

- We propose an efficient method to compare the predicted perceptual discriminabilities of two models based on their metric tensors.
- We apply the comparison method to two neural models for human color perception. We used the comparison results to synthesize a hybrid model that outperforms both existing models on predicting human chromatic discrimination [11].
- We apply the method to two neural network architectures (autoencoders trained with L1 or L2 regularizers), and showed that the former better matches human perception.

## 2 Methods

### 2.1 Computing metric tensors for deterministic and stochastic neural models

#### Metric tensors for deterministic representations

Suppose an image **s** consists of *p* pixels 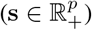. A deterministic neural encoding model maps the image **s** to *n* neurons’ mean response rates 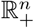. Existing comparison methods typically examine neural representations for a discrete set of images,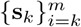(like the middle panels in Fig. 1). We take a slightly different approach, by assuming that the model smoothly and *differentiably* varies with the stimulus, and we examine the differential change of the model predictions due to stimulus perturbations.

We use ***ϵ***∈ ℝ^*p*^ to denote a perturbation to image **s**. ∥***ϵ***∥ is assumed small, so that when the image is perturbed, the neural response change can be linearly approximated (via Taylor expansion) as the derivative of ***µ***(**s**) along the ***ϵ*** direction, which we denote as **d**_***ϵ***_(**s**). **d**_***ϵ***_(**s**) can be expressed as a matrix product (a linear computation) between the Jacobian of the neural response, *J*_***µ***_(**s**), and the perturbation ***ϵ***: **d**_***ϵ***_(**s**) = *J*_***µ***_(**s**)***ϵ***. Further, we assume that *perceptual discriminability p*_***ϵ***_(**s**) captures the *L*_2_ norm of the neural response change, which has been supported by empirical evidence ([1, 2]):

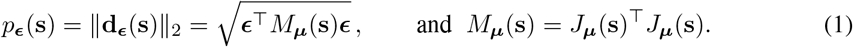

Here, *M*_***µ***_(**s**) is a metric tensor. It is a *p* × *p* matrix that is symmetric and positive definite (assuming it is full-rank for now), and it determines the model’s discriminability prediction along arbitrary perturbation direction ***ϵ*** to image **s**. *M*_***µ***_(**s**) completely summarizes the distance structure of the neural representation around ***µ***(**s**). Geometrically, *M*_***µ***_(**s**) transforms the equi-length image perturbations ***ϵ*** (a *p*-dimensional ball) into a *p*-dimensional ellipsoid. The predicted discriminability is the largest along the largest eigenvalue direction of *M*_***µ***_(**s**) (the major axis of the ellipsoid) [12]. Because neural encoding models are generally nonlinear, for two distinct reference images **s**_1_ and **s**_2_, *M*_***µ***_(**s**_1_) and *M*_***µ***_(**s**_2_) are generally different.

#### Metric tensors for stochastic representations

A stochastic representation ***ν*** maps an image **s** to an *n*-dimensional neural response distribution *P* (**r** | **s**). We assume that ***ν*** is differentiable in the sense that a well-defined Jacobian for the log of the density function *P* (**r** | **s**) exists, which we denote as *J*_log ***ν***_(**s**). Fisher information is the stochastic analog of the metric tensor *M*_***µ***_(**s**), and can be expressed as the expectation of a quadratic form of the Jacobian (see S2):

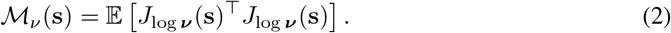

Fisher information has been used to summarize discriminability in recent perceptual literature [13, 14, 15, 16, 17, 18]. Computing Fisher information is generally non-trivial for arbitrary distributions, and experimentally, neural response distributions are rarely verified beyond the first two moments. In practice, a lower bound on Fisher information [19] can be used to summarize perceptual discriminability [20, 3]. This lower bound computation only involves the first two moments of the neural response distribution *P* (**r**|**s**). We use *M*_***ν***_(**s**) to denote Fisher lower bound, or the metric tensor of interest for stochastic neural models. To be consistent with the notations for deterministic models, we use ***µ***(**s**) to denote the mean neural response as a function of stimulus, and Σ(**s**) as the response covariance matrix, and perceptual discriminability can be expressed analogously to that of Equation 1:

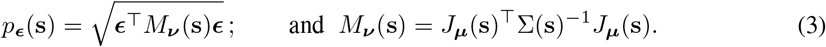

Notice that in addition to the Jacobian of the mean neural response, the metric tensor for stochastic neural models also depends on the response covariance matrix.

#### Perceptual discriminability and thresholds

To compare between two neural models **f**_1_(**s**) and **f**_2_(**s**), we can compare their metric tensor predictions *M*_1_(**s**) and *M*_2_(**s**). We can apply metric tensor comparison to deterministic or stochastic models, or a mixture of the two, so the method is more flexible than existing model comparison methods. For simplicity of visualization, as well as being consistent with the perceptual literature, in the rest of the paper, we examine the threshold matrix *T*_**f**_ (**s**), the inverse of discriminability:

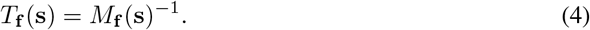

Like *M*_**f**_ (**s**), *T*_**f**_ (**s**) is also symmetric and positive definite (it is also a metric tensor). Expressing results in terms of thresholds *T*_**f**_ (**s**) has the advantage that the values share the same units as the stimuli, and can be visualized in the stimulus space.

### 2.2 Comparing metric tensors at a single reference image

Different neural models **f**_1_ and **f**_2_ generally make different metric tensor predictions (*T*_1_(**s**) and *T*_2_(**s**)) at a reference image. To assess which of the two model predictions is more similar to human perceptual thresholds measured at **s**, denoted as *T*_**h**_(**s**), a direct approach would be to compare matrix distance for the pair {*T*_1_(**s**), *T*_**h**_(**s**)}, and the pair { *T*_2_(**s**), *T*_**h**_(**s**) } using some matrix distance measure [21] (e.g. Frobenius norm). The model prediction that is closer in distance to human perceptual thresholds is the preferred model. Empirical estimation of human perceptual thresholds *T*_**h**_(**s**) is infeasible in a high-dimensional space. Suppose human thresholds *T*_**h**_(**s**) have dimensionality *r* (*r*≤*p*), and suppose the sampling noise is bounded (and assuming i.i.d. sub-Gaussian samples of ***ϵ***), stable recovery of *T*_**h**_(**s**) takes an order of *pr* samples [22, 23]. If an image **s** consists of 100× 100 pixels, and *T*_**h**_(**s**) is assumed to have rank 30 (a conservative estimate), measuring threshold along each ***ϵ*** perturbation takes around 2 minutes, and the total amount of time to recover *T*_**h**_(**s**) is in the order of 10,000 hours per human subject. The direct approach of model comparison is not experimentally feasible, so we introduce an alternative model comparison method, which is much more efficient, and requires as little as a *a single trial* to distinguish between two metric tensor predictions.

#### Full-rank metric tensor comparisons

We first make an observation that making perceptual threshold measurements using some image perturbations are more informative to distinguish between metric tensors than the others. Examine Figure 2A, measuring the human perceptual threshold along perturbation ***ϵ***_*B*_ would not distinguish between the two models, because the two models’ predictions along ***ϵ***_*B*_ are identical. On the other hand, the two metric tensors make very different predictions along perturbation ***ϵ***_*A*_. To efficiently sample human perceptual thresholds to distinguish between model-predicted metric tensors, we need to find image perturbations that the metric tensors make *the most distinct predictions*.

**Figure 2:**
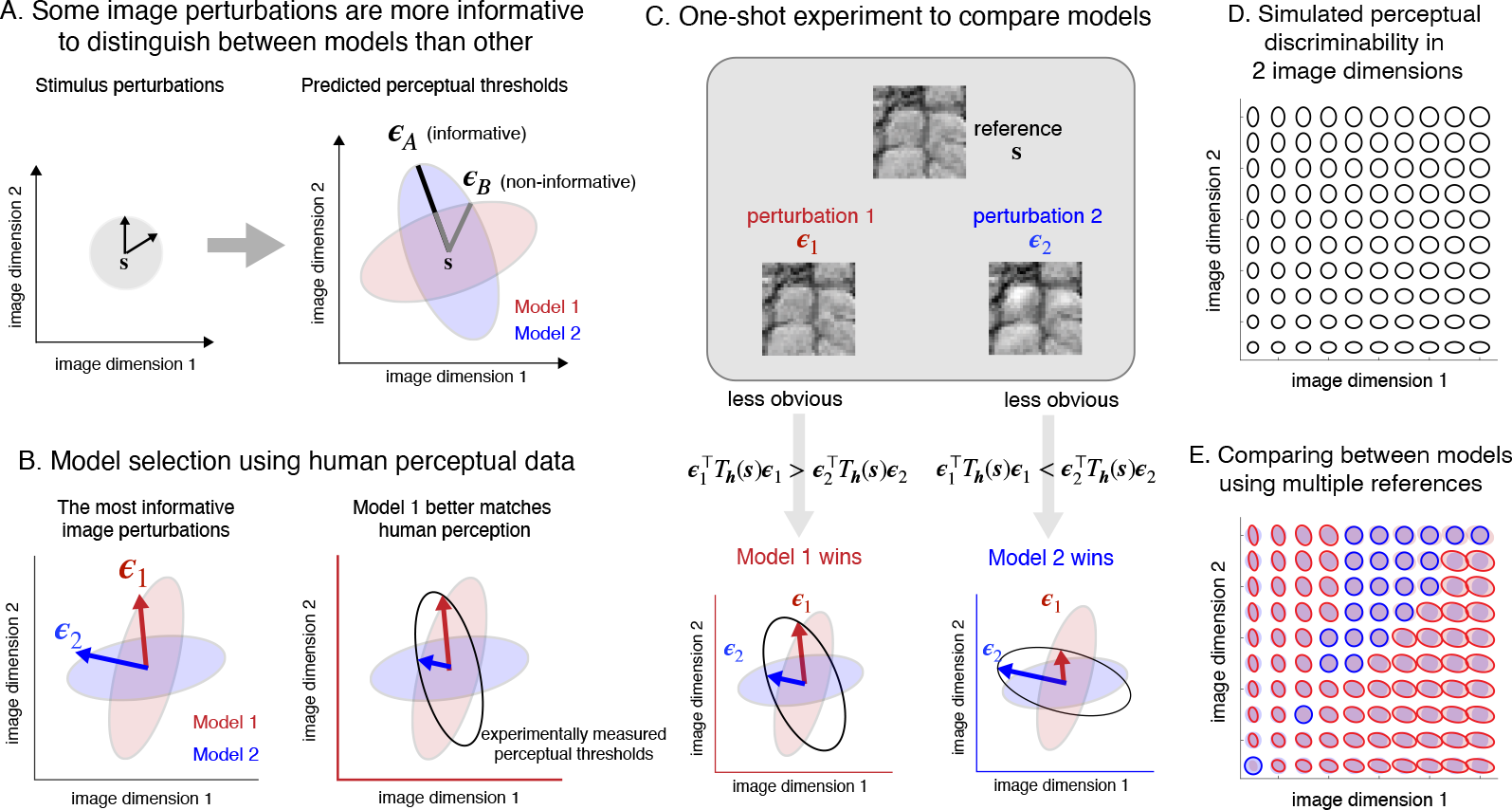
Model comparison using metric tensors. **A**. Locally, a neural model transforms a circle of equidistance perturbations of a reference image **s** to a two-dimensional ellipse. Different models generally predict different ellipses (hence different metric tensors). Not all perturbation directions are equally informative to distinguish the two metric tensors. For this example, The two tensors predict distinct thresholds along image perturbation ***ϵ***_*A*_, but identical thresholds along image perturbation ***ϵ***_*B*_ . **B**. We compute (see text) perturbations that maximally distinguish between the two metric tensors’ threshold predictions. To compare the models to human perceptual discriminability, we only need to measure perceptual discriminability along the two chosen perturbation directions. **C**. Perceptual experiment for model selection. An observer is asked which of the two perturbed images is more similar to the reference. If perceptual threshold is larger along ***ϵ***_1_ (as compared to ***ϵ***_2_), model 1 wins, and vice versa. **D**. Simulated human metric tensors in a two-dimensional stimulus space at uniformly sampled references. **E**. We simulated two models with different metric tensor predictions at different references in the two-dimensional image space. The color of an ellipse indicates the winning model for that reference image. Overall, model 1 makes metric tensor predictions more similar to human discriminability, but model 2 out-performed model 1 in some region of the stimulus space.

To do so, we find an ***ϵ*** that maximizes the ratio between the two metric tensor predictions. Because one model’s prediction can be much larger in amplitude than the other, and we want to find the metric tensor that is more similar to human perceptual thresholds in shape (or similar in terms of relative thresholds in different directions). To eliminate the effect of predicted metric tensor size (see S4), we seek a pair of image perturbations ***ϵ***_1_ and ***ϵ***_2_, such that ***ϵ***_1_ maximizes the ratio between the metric tensor predictions, and ***ϵ*** maximizes the inverse of the ratio:

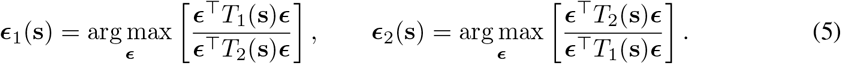

To find the ***ϵ***_1_ and the ***ϵ***_2_ is to solve two generalized eigenvalue problems, which have well-known closed-form solutions: ***ϵ***_1_ is the eigenvector that corresponds to the largest eigenvalue of matrix *T*_2_(**s**)^−1^*T*_1_(**s**); symmetrically, ***ϵ***_2_ is the eigenvector that corresponds to the largest eigenvalue of *T*_1_(**s**)^−1^*T*_2_(**s**). We assumed both matrix *T*_1_ and *T*_2_ are full-rank for now, so we can compute their inverse, and later on we consider the case when either matrix (or both) are rank-deficient. In the first panel of Figure 2B, we illustrate the optimal pair of image perturbations chosen for two simulated metric tensors.

Once a pair of image perturbations is chosen, we need to make human perceptual measurements along these perturbations. One advantage of our method is that to choose a winning model, we do not need to measure the absolute perceptual thresholds along the two perturbation directions, but we only to assess the relative thresholds – which of the two perturbations is more perceptually salient? For example, in the second panel of Figure 2B, if perceptual threshold measured using perturbation ***ϵ***_1_ is greater than that using ***ϵ***_2_, then we conclude that model 1 is closer to predicting the human thresholds than model 2. We use *t*_*i*_(**s**) to indicate perceptual threshold measured along ***ϵ***_*i*_, and to experimentally distinguish between the two models, we only need to determine whether *t*_1_(**s**) *> t*_2_(**s**), or *t*_2_(**s**) *> t*_1_(**s**), and equality only occurs when the two metric tensors are identical up to a scale factor.

During a model-selection experiment (Figure 2C), an observer is presented with a reference image **s**, together with two perturbed versions of the image (**s** + ***ϵ***_1_) and (**s** + ***ϵ***_2_). The observer judges which of the two perturbations is more noticeable. We use≻ to indicate model preference, and the model comparison can be summarized as:

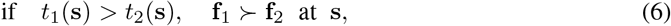

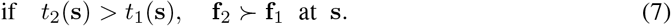

Notice that larger perceptual threshold (or smaller perceptual discriminability) means the image perturbation is less noticeable.

#### Rank-deficient metric tensor comparisons

A metric tensor can be rank-deficient when the number of linearly independent neurons is less than the number of image pixels. In this section, we conduct analyses analogous to the full-rank case, but using model-predicted discriminability *M*_**f**_ (**s**) instead of threshold *T*_**f**_ (**s**), because *M*_**f**_ (**s**) can be directly derived from a neural model, and is no longer assumed invertible. We introduced two different approaches to extend our model comparison method to rank-deficient metric tensors, and we will use each of the approaches in one of our applications.

In the first approach, we reduce the stimulus dimensionality via a map 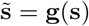, and 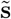can be either linear (e.g. cropping an image) or nonlinear (e.g. only examine perceptual discrimination by varying image contrast). We assume that neurons take inputs from the transformed stimulus 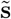, and map into the response space via **f** [**g**(**s**)], and the corresponding metric tensor *M*_**f**_ 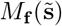(for the computation from the transformed input space), relates to the metric tensor *M*_**f**_ 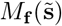 for the neural computation from the original stimulus space via:

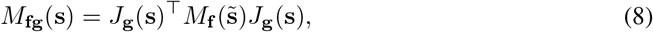

and *J*_**g**_(**s**) is the Jacobian for the image transform **g**(**s**) (see S5 for details).

In the second approach, we modified our objective functions (Equation 5) to accommodate the rank deficiency in *M*_*i*_(**s**). Intuitively, we search for an image perturbation that maximizes the numerator computation within the denominator metric tensor’s null space. We useV_*i*_ to denote the nullspace of *M*_*i*_(**s**), and the modified objective functions can be expressed as:

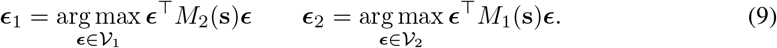

This approach is conceptually similar to Equation 5, and see S6 for more computational details.

### 2.3 Comparing metric tensors using multiple reference images

Comparing metric tensors using a single reference image is informative. But to achieve a more general understanding of how each model captures human perceptual representations, we may compare metric tensors measured using multiple (or a large set of) reference images 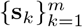 . To select between two models across multiple reference image, we first compare model-predicted metric tensors for each reference **s**_*k*_. Let **1**_*T*_*i* (**s**_*k*_) be an indicator function which is 1 when *T*_*i*_(**s**) is the winning metric tensor at **s**_*k*_, and 0 otherwise. We further use *c* [*T*_*i*_] to denote the sum of the indicators for the *i*^*th*^ model across all references. The winning model satisfies:

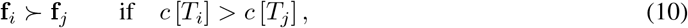

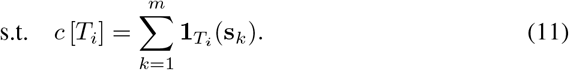

Using our model comparison method, we not only obtain a winning model, but also obtain a lot of additional information about human representation through the comparison. For example, in Figure 2DE, we compared between two neural models using uniformly sampled references in a two-dimensional image space. Comparing model-predicted metric tensors across references to simulated human perceptual measurements, we observe that one model better captures perceptual discriminability in a certain part of the image space, while the other model better predicts the rest of the space. The fusion of these models, can possibly yield an improved model of human perceptual discriminability for the entire image space.

## 3 Application

### 3.1 Simple color discrimination example

We first examine a toy example in color discrimination. Within an equi-luminant two-dimensional color space (also called *xy* space, or chromatic space), we choose a reference color – a neutral gray, and showed six equi-distance perturbations of the reference (Fig. 3A). All equi-distance perturbations form a circle in the two-dimensional chromatic space. We simulated two models that make distinct chromatic discriminability predictions. Both models’ predictions transform the circle of equi-distance perturbation to ellipses, but the two models predict ellipses that differ in their major and minor axes. Human perceptual thresholds measured around the reference image also exhibit an elliptical shape (Fig. 3B). To examine which model’s threshold predictions are closer to those of humans, we found two effective image perturbation directions using the method described in the previous section (Fig. 3C). With the effective perturbation directions, we can conduct a human experiment to test the relative perceptual thresholds along the two directions. If perturbation 1 in Fig. 3D is viewed as less obvious compared to perturbation 2, then model 1 is the winning model, and vice versa.

**Figure 3:**
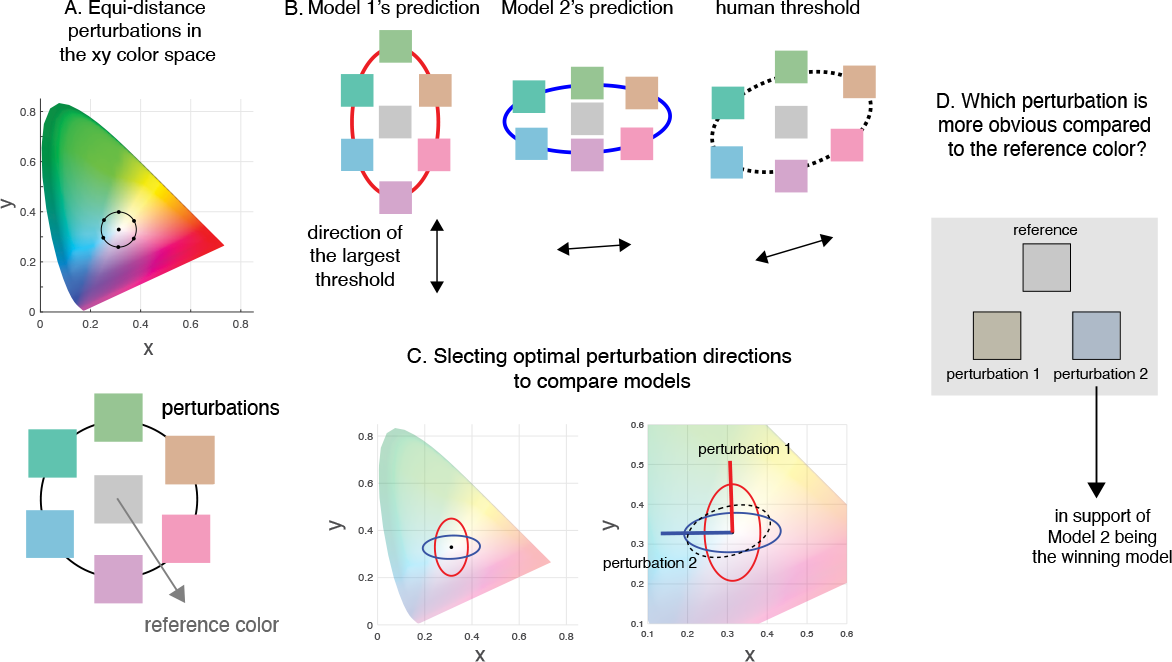
A toy example. **A**. A reference color (a neutral gray) was perturbed along six different directions, and the equi-distance perturbations form a circle in the 2D chromatic space. **B**. Two simulated models, as well as human perception, transform the circle of stimulus perturbation into three different two-dimensional ellipses in the 2D chromatic space. **C**. We choose two efficient perturbation directions to determine which model better matches human perceptual thresholds around the reference. **D**. A perceptual experiment can be conducted to compare relative thresholds along the two efficient perturbation directions, determining which model is best aligned with human perception.

### 3.2 Application 1: Comparing neural encoding model for chromaticity

Here, we examine a more realistic perceptual example to illustrate how our method differentiates between models, selects the one that better matches human perceptual discriminability, and can aid generating improved neural encoding models. A chromaticity diagram describes a two-dimensional color space that varies in hue and saturation (with fixed luminance, and restricted to human range of visibility). MacAdam measured two-dimensional perceptual thresholds within the chromaticity diagram (Figure 4A,[11]), a heroic effort requiring 25,000 experimental trials for each observer. A recently proposed neural encoding model [17] provides a reasonable prediction of the seemingly intricate chromatic threshold patterns (Fig. 4B). In the proposed model, three cone types (*L, M*, and *S* cones) linearly respond to incoming light, and each cone’s linear output is paired with a noise model. The noise model can either be independent Poisson-distributed [17], motivated by photon noise in the incoming light, or it can be independent Gaussian-distributed, motivated by noise properties of cone photoreceptor responses (Figure 4BC, see S7 for computational details).

**Figure 4:**
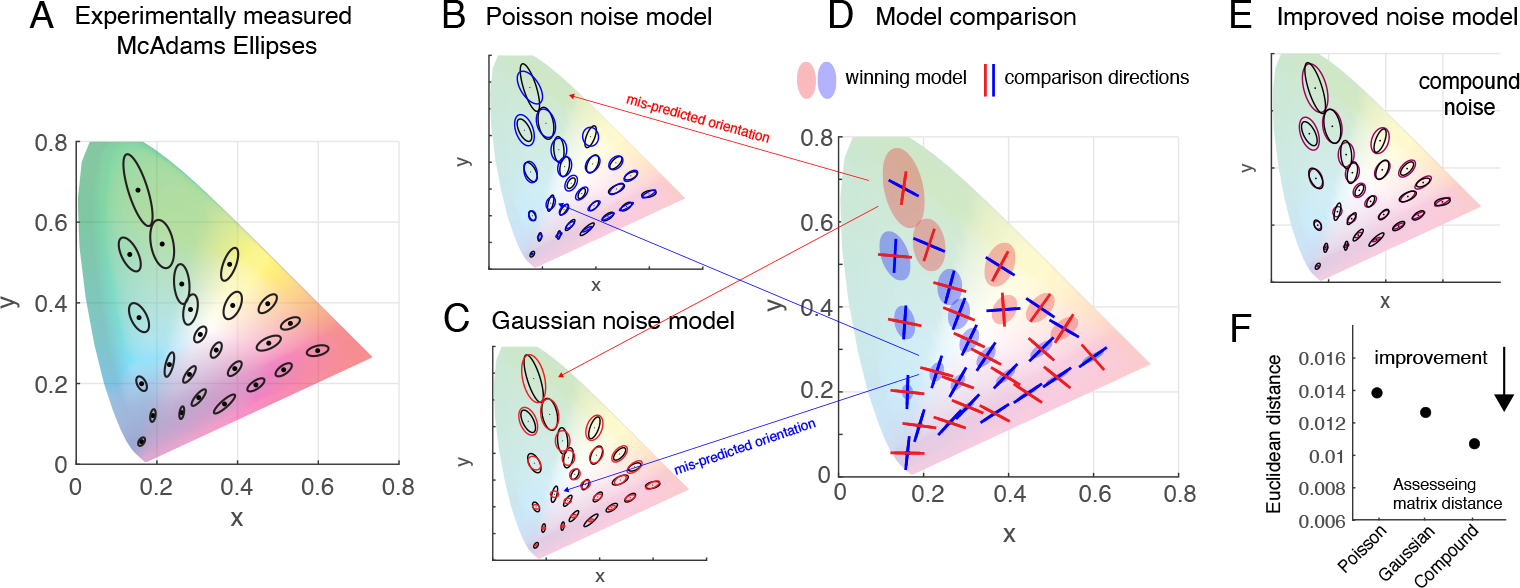
Comparing neural encoding models for chromatic thresholds. **A**. Experimentally measured chromatic thresholds by MacAdam [11]. The two-dimensional ellipses illustrate the measured threshold, which are super-imposed on the translucent chromatic diagram in the *XY* color space. **B**. Predicted chromatic thresholds using linear cone model combined with independent Poisson noise [17]. **C**. Predicted thresholds using linear cone model combined with independent Gaussian noise. **D**. Model comparison. We illustrated the two optimal comparison directions for the model pair at each reference, and the color of the ellipse underneath illustrates the winning model at that reference. **E**. Predicted chromatic thresholds using the compound noise model. **F**. The compound model improves upon both the Poisson and the Gaussian model in terms of Euclidean matrix distance (see S8 for computational details).

To select between these two noise models, we applied our model comparison method at each reference color, and found two optimal perturbation directions per reference color. If we were to perform model selection on a new human observer, only 25 trials (approximately 3 minutes of experimental time) are required to differentiate between the two models. Thus, our model comparison method offers a means to bridge the gap between the high-dimensional representations examined by the neuroscience and machine learning communities, and the low (often one-) dimensional measurement methodologies of the perception community.

The predicted chromatic metric tensors using both models reasonably resemble the chromatic thresholds measured by MacAdam. But closer inspection reveals that the Poisson noise model tends to mis-predict metric tensor orientations in the green-yellow region, and the Gaussian noise model tends to mis-predict the orientations in the purple-red region. This suggests that a compound model can potentially outperform each noise model alone. To this end, we propose the following noise model (for the *i*^*th*^ cone type):

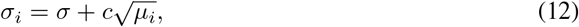

where *σ* is a constant (corresponds to Gaussian noise), and *c* is a non-negative scalar (corresponds to Poisson-like noise). We find that this compound model out-performed both existing noise models, and we illustrate the compound model and the comparison of model performance in Fig. 4E and 4F.

### 3.3 Application 2: Comparing autoencoders trained with *L*_1_ or *L*_2_ regularizers

In this application, we trained two autoencoders with a single hidden layer. The weights of the first auto-encoder were trained with an L1 regularizer (to encourage sparsity), and the weights of the second auto-encoder were trained with an L2 regularizer (to encourage smoothness between weights). The auto-encoders were trained on MNIST, and the reconstruction errors of the autoencoders were qualitatively similar: 1.7e-2 for L1 and 1.2e-2 for L2 on average across all images. After training using stochastic gradient descent on the weights and biases, we computed the Jacobian of the hidden layer representation for a reference image, which was then used to compute the metric tensor. Because of the regularizers, metric tensors derived from the models are generally low rank, so we used Eq. (9) as objective to find the optimized image perturbation directions for each reference.

For each of the 200 MNIST images (references), we found two most efficient perturbations to distinguish between the two auto-encoders’ predicted perceptual discriminability. Notice that in the previous application, the model-predicted discriminability at one reference image is a two-dimensional ellipse, and in this application, the predicted discriminability is a high-dimensional ellipsoid. Using the experimental method described in Fig. 3, we tested relative discriminability of the two model-derived perturbations for 200 MNIST digit images. We find that the L1-trained autoencoder is better aligned with human perception for most images: The participants selected L1 model as preferred model for 78% (participant 1), and 80% (participant 2). In Fig. 5, we demonstrate the perceptual experiments using two reference images as examples.

**Figure 5:**
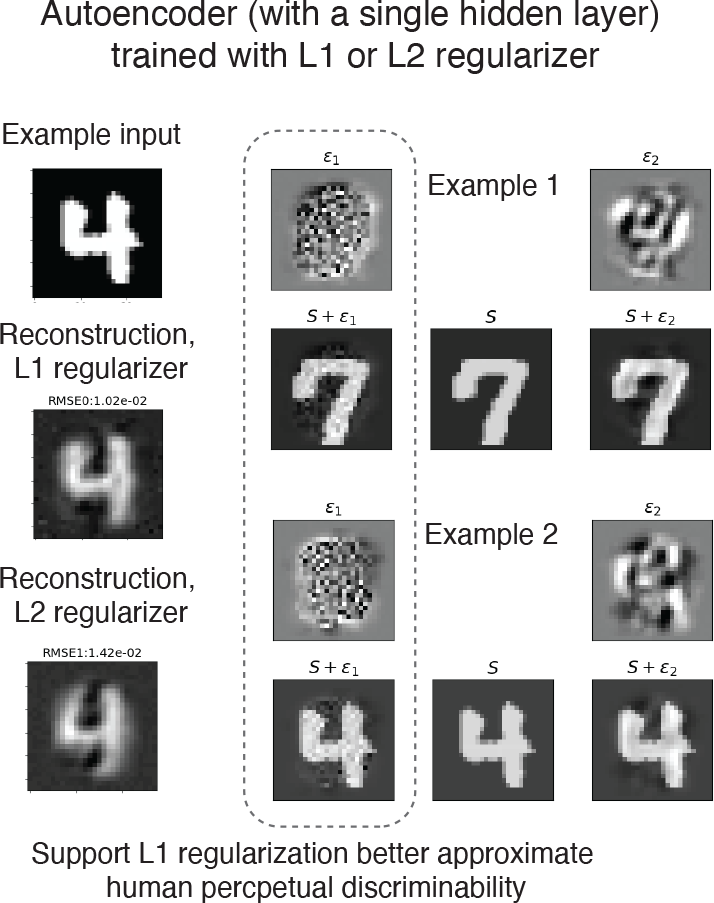
Comparing metric tensors of autoencoders trained using L1 or L2 regularizer. First, we demonstrate two reconstruction examples, one trained with L1 regularizer, and the other trained with L2 regularizer. The reconstruction errors across the two types of models are qualitatively similar. Then we demonstrate two sets of perceptual experiments, which involves a reference image, and two image perturbations. The perturbation on the left to the reference image seem more perceptually salient to participants, indicating that the autoencoder trained with L1 regularizer better captures human perceptual discriminability for this image.

## 4 Discussion

We have proposed a novel methodology for comparing two models in terms of their perceptual discriminability predictions. The method is flexible – it can be applied to either deterministic or stochastic models, or a mixture of two. It is also extremely efficient – for a given reference stimulus, a human experiment testing a single pair of perturbed stimuli is sufficient to determine which of two models best accounts for human discriminability around that reference.

Our model comparison method is a generalization of the “Eigen-distortion” method [12], which generates image perturbation that maximizes (or minimizes) a single model’s predicted perceptual discriminability. Beradino et al. find the minimal/maximal solutions of the Rayleigh quotient problem, whereas our method finds the minimal/maximal solutions of *generalized Rayleigh quotients*. The eigendistortions can be used to provide an indirect comparison of models (see [12]), but note that two models can be identical in their maximal/minimal perturbation directions [12], but still be distinguishable using our method.

Our method is based on perturbations that maximize/minimize the *ratio* between two metric tensor predictions. Alternatively, the problem can be formulated using other contrastive objectives, such as maximizing the *difference* between the two metric tensor predictions. The difference objective can also be solved with closed-form solutions (see S9). But unlike the ratio objective, for which re-scaling one metric tensor does not change the optimal image perturbation directions, the difference objective (and resuting perturbation directions) is affected by re-rescaling.

Our method can be generalized to compare more than two models simultaneously, while maintaining its advantages in terms of experimental efficiency. Instead of comparing the ratio of two metric tensor predictions, we can compare the ratio between a metric tensor’s prediction, and the average of all models’ metric tensor predictions, and the problem remains equivalent to a generalized eigenvalue problem (see S10). We’ve not yet tested and validated this generalized method, but it offers the possibility of additionalincreases in the efficiency of comparing model predictions to human perception.

